# Prevalent pH Controls the Capacity of *Galdieria Maxima* to Use Ammonia and Nitrate as a Nitrogen Source

**DOI:** 10.1101/785287

**Authors:** M. Iovinella, DA. Carbone, D. Cioppa, S.J. Davis, M. Innangi, S. Esposito, C. Ciniglia

**Affiliations:** Department of Biology, University of York, YO105DD York UK; Department of Biology, University of Naples Federico II, 80126 Naples Italy; Department of Environmental, Biological and Pharmaceutical Science and Technology, University of Campania “L. Vanvitelli”, Caserta, Italy

**Keywords:** *Galdieria*, pH, Ammonium, Nitrate

## Abstract

*Galdieria maxima* is a polyextremophilic alga capable of diverse metabolic processes. Ammonia is widely used in culture media typical of laboratory growth. Recent reports that this species can grow on wastes promote the concept that *G. maxima* might have biotechnological utility. Accordingly, there is a need to know the range of pH levels that can support *G. maxima* growth in a given nitrogen source. Here, we examined the combined effect of pH and nitrate/ammonium source on the growth and long-term response of the photochemical process to a pH gradient in different *G. maxima* strains. All were able to use differing nitrogen sources, despite both the growth rate and photochemical activity were significantly affected by the combination with the pH. All strains acidified the NH_4_^+-^medium (pH<3); only *G. maxima* IPPAS P507 showed reduced capacity in lowering the pH from 6.5. pH was a limiting factor in nitrate uptake at pH`≥`6.5; noteworthy, at pH 5 on nitrate *G. maxima* ACUF551 showed a good growth performance, despite the alkalization of the medium.

## 1. INTRODUCTION

Polyextremophiles are organisms able to survive and grow under more than one harsh environmental conditions, such as extremes of temperature (thermophily), pH (acidophily, alkaliphyles), salts (halophily), and pressure (barophily) [1, 2]. The ability to counteract environmental stressors is related to the development of specific mechanisms concerning structural and biochemical adaptation, gene expression and regulation. These make such organisms essential sources for biotechnological applications [1].

Geothermal environments, characterized by high temperatures (above 50°C), low pH values (<1), and high amounts of heavy metals, are common areas where polyextremophiles are undisputed rulers: bacteria, fungi and some algae are the only surviving microorganisms [3, 4]. Among the latter, Cyanidiophyceae (Rhodophyta) represent almost all the thermoacidophilic eukaryotic biomass [5]. Cyanidiophyceae (*Galdieria sulphuraria*, *G. maxima*, *G. phlegrea, Cyanidium caldarium*, and *Cyanidioschyzon merolae*) are the most abundant photosynthetic protists found in extremely acidic, sulfur-rich environments generally bordering active volcanoes [6–14]. Condensation of sulfur dioxide and hydrogen sulfide produces sulfur crystals, which are subsequently oxidized to sulfuric acid resulting in acidification (pH 0.5-3). Sulfur allows cells to produce S-containing molecules with high antioxidant power, such as glutathione [3,15,16].

Recent genome sequencing demonstrated that horizontal gene transfer (HGT) from prokaryotic sources provided *G. sulphuraria* extraordinary metabolic flexibility. It is able to use more than 50 different carbon sources [17–21] and to survive in hostile environments rich in heavy and rare metals (e.g., genes to detoxify mercury and arsenic). The extent of metabolic flexibility for other elements, such as prevailing N is less explored, but it is known that amino acids can be used as an N-growth component [22].

So far, Cyanidia were isolated from several solfataras worldwide (Italy, Iceland, Japan, New Zealand, Yellowstone National Park, and Taiwan). The acidophily/acidotolerance was considered as one of the main factors limiting the dispersal and survival of Cyanidiophyceae; nevertheless, recent explorations in Turkey provided the ubiquity of cyanidia in several Turkish thermal baths, mostly on neutral and sub neutral soils (pH 5.8-7), growing as thin biofilms hypolithically and endolithically [13]. The ability to lower the external pH from 6 to more acidic values was tested by Lowell and Castenholtz (2013) on several Cyanidiophiceae strains collected from Yellowstone, Japan, Philippines, and New Zealand hot springs, suggesting the importance of this process as a survival strategy in nonacidic environments during long-distance migration [3,20,23]. Growth in the sub-neutral medium was also evaluated by Henkanatte-Gedera et al. (2017), both at laboratory and field scale. These data confirmed the capacity of *G. sulphuraria* to grow in a broader range of initial environmental pHs, suggesting the potential savings in chemical costs associated with pH adjustment of real waste streams [24].

The employment of *Galdieria* species in applied sciences prompted us to assess the effect of initial pH on the growth of eight *G. maxima* strains collected from acidic, neutral, and sub-neutral thermal areas. Likewise, since proton exchange can be dependent on the uptake and metabolism of nitrogen compounds [25], we investigated the combined effect of pH and nitrate/ammonium source on their growth. We also evaluated the long-term response of the photochemical process to a pH gradient, considering the maximum efficiency of photosystem II (Fv/Fm) and the non-photochemical quenching (NPQ) as indicators of the physiological state of the cultures as a response to pH and Nitrogen source.

## 2. MATERIAL AND METHODS

### 2.1 Strains cultivation

Eight *Galdieria maxima* strains, supplied from the Algal Collection of University Federico II of Naples (www.acuf.net) and formerly collected from Turkey [13], Iceland [9, 12], and Russia [16], were selected based on the pH of the original collection site (Table 1). All strains were inoculated in Allen medium containing (NH_4_)_2_SO_4_ as the nitrogen source, at pH 1.5 [26] and cultivated at 42°C under continuous fluorescent illumination of 50µmol photons·m^-2^·sec^-1^ until they reached the exponential phase.

**Tab. 1.** Code, name of the collection sites, environmental pH and temperature

| Strains | Sampling site | Temperature (°C) | pH |
| --- | --- | --- | --- |
| IPPAS_P507 | Kunashir (RUS) | 41.5°C | 4 |
| ACUF769 | Güçlükönak (TURK) | 60°C | 1 |
| ACUF722 | Güçlükönak (TURK) | 60°C | 1 |
| CloneT18 | Diyadin (TURK) | 45°C | 7 |
| ACUF773 | Diyadin (TURK) | 45°C | 7 |
| ACUF648 | Kula, Manisa (TURK) | 41°C | 7 |
| ACUF731 | Kula, Manisa (TURK) | 41°C | 7 |
| ACUF551 | Landmannalaugar, Hekla (ICE) | 42°C | 1 |

### 2.2 Phylogenetic analysis

We retrieved from GenBank 197 rbcL nucleotide sequences from different Cyanidiophyceae species and strains [6–9,12,13]; DNA extraction from *G. maxima* ACUF551 was performed using the protocol described in Ciniglia et al., 2015 [27]. Four degenerate primers were used to amplify the rbcL gene from the sample [6]. The resultant product was purified with the QIAquick PCR purification kit (Qiagen) and used for direct sequencing using the BigDyeTM Terminator Cycle Sequencing Kit 3.1 (PE-Applied Biosystems, Norwalk, CT, USA) and an ABI-3500 XL at the Microgem Laboratory (Naples, Italy). Forward and reverse electropherograms were assembled and edited using the program Chromas Lite v.2.1 (http://www.technelsium.com.au/chrom as.html). All sequences were aligned with the MUSCLE command in UGENE v 1.16.0 [28, 29]; four species belonging to the subphylum Rhodellophytina (*Rhodella violacea*, *Dixoniella grisea*, *Porphyridium aerugineum*, *Bangiopsis subsimplex*) Rhodophyta, were chosen as outgroup (Table S1). No gaps or indels were incorporated in the alignment, and the best likelihood tree was estimated under the general time reversible (GTR) substitution with gamma-distributed rate heterogeneity (G) and proportion of invariable sites estimation (I) as result of model-testing in jModeltest v.2.1.10 [30]. Maximum likelihood (ML) analysis were performed with RAxML v.8.2.8 using the rapid bootstrap option (1000 replicates) to evaluate the branches statistical support [31].

### 2.3 Experimental procedure

Microalgal cultures (OD∼0.35; 1.5-2.0•10^7^ cells) were inoculated in fresh Allen medium under different pH levels (pH 5, 6, 6.5 and 7) to investigate the influence of pH on growth; control tests were performed at pH 1.5. These experiments were carried out in 50ml Erlenmeyer flasks containing Allen medium with (NH_4_)_2_SO_4_ as the nitrogen source (10mM) at 42°C under continuous cool fluorescent light (50 µmol photons·m-2·second-1), and kept mixed on an orbital shaker illuminated beneath. All analyses were performed in triplicate.

The pH of the medium was measured weekly for six weeks (Mettler, Toledo) and the growth rates were monitored by spectrophotometric measurements of the optical density OD_550_ (Secomam spectrophotometer Prim light). The effect of the pH on the growth of the algal strains was assessed by calculating the maximum growth rate using the equation:

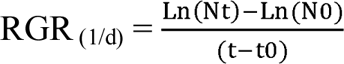

Where: Nt is the optical density at the final time

N_0_ is the optical density at the initial time

T is the final time (days)

T_0_ is the initial time (days)

The effect of different nitrogen source was next evaluated on three strains (IPPAS P507, ACUF722 and ACUF551) grown in Allen medium where the ammonium was replaced by NaNO_3_ in an equimolar quantity, at pH 7, 6.5, 5, and 1.5, using the same experimental design of previous experiments. In these assays, tests were also performed in Allen medium containing ammonium at the same pHs. Chlorophyll a content (AquaFluorTM; Handheld Fluorometer/Turbidimeter; Turner Designs) and pH variations were recorded at 4-days intervals during the experiments for 12 days. All measurements were taken in triplicates.

Photochemical activity of IPPAS P507, ACUF722, and ACUF551 was determined with Pam Fluorometry [32] at 42°C in a thermostatic chamber (Hansatch) by a pulse amplitude modulated fluorometry (Hansatech fluorescence monitoring system), following Hanlet protocol for red algae [33]. After a dark period (30 min), 2 ml of each sample were placed in a quartz cuvette near the optic fiber of fluorimeter, under continuous stirring. During the analyses, microalgae were exposed at four different light intensity (31μmol photons m^−2^s^−1^, 65 μmol photons m^−2^s^−1^, 116 μmol photons m^−2^s^−1^, and 281 μmol photons m^−2^ s^−1^), μ alternating three minutes of light and three minutes of dark [34]. Three variables were considered, the maximum efficiency of photosystem II, non-photochemical quenching and quantum yield of photosystem II. The maximum efficiency of photosystem II (Fv/Fm), index of the relatively good physiological state of the cultures, was evaluated in the dark with a saturating light pulse of 2900 μmol photons m^−2^s^−1^ at 0.6s. Non-photochemical quenching (NPQ) was determined by the formula (Fm – F0)/F0 and used as an indication of algal stress condition [35]. The quantum yield of photosystem II (FPSII) is an indicator of PSII photochemical efficiency in the presence of light. All measurements were done in triplicates.

### 2.4 Statistical analysis

All experiments were carried out in triplicate and results were reported as mean ± standard deviation. Variables were inspected for normality and homogeneity of variance. Variations of the relative growth rate, culture medium pH, chlorophyll content, and photochemical activity were tested by means of one-way analyses of variance (one-way ANOVA). As multiple comparison post-hoc test to evaluate the significance of differences among treatments, Tukey HSD was used (α = 0.05). All statistical analyses were performed with R studio software [36].

## 3. RESULTS

### 3.1 Phylogenetic analysis

Molecular characterization, based on the partial (842 bp) rbcL phylogeny, classified all the isolates chosen for these experiments as *G. maxima* (Fig. 1). The analysis identified a divergence of *G. maxima* in two subclades (100% MLB), and the grouping of most of Turkish *G. maxima* isolates in the sub-population GmA (100% MLB). All these isolates shared 100% of identical sites with the Russian strain IPPAS P507 from Kamchatka peninsula. The other subclade (GmB) comprised other Turkish strains and the ones collected from the Icelandic areas (74% MLB). The rbcL sequence of the Icelandic isolate ACUF551, belonging to this group, slightly diverged from the reference sequence (IPPAS P507) sharing with it 95.6% of identical sites.

**Fig. 1.**
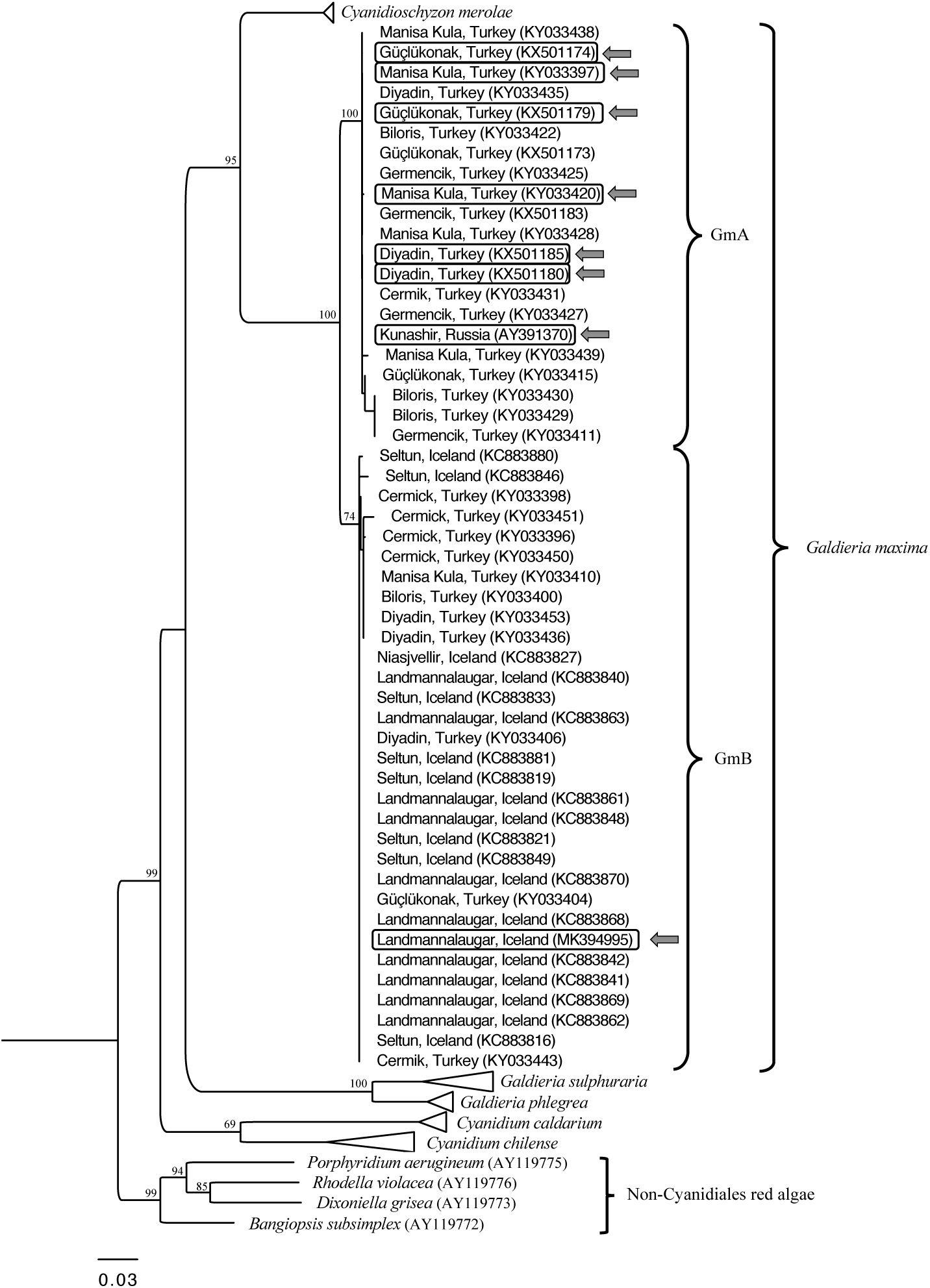
Maximum likelihood tree of Cyanidiophyceae analysis based on rbcL sequences. Only bootstrap values > 60% are shown near nodes. The arrows indicate the *G. maxima* strains selected for the experiments. See Tab. S1 for details.

### 3.2 Relationship between pH and growth under ammonium source

The growth of 8 *G. maxima* strains was followed and related to the lowering of pH for six weeks, starting from different pHs and providing NH_4_^+^ as the nitrogen source. At an initial pH 1.5, the external pH remained stable, with low and non-significant variations among all strains and throughout the experiments. In contrast, when considering a starting pH 7, none of the strains displayed detectable growth after the six weeks, and small variations of pH were recorded in all cultures (Table S1).

Plotting the concurrent lowering of external pH with the increase of biomass yield during the experiments started from pH 5, 6, and 6.5, a correlation existed between the final pH and the biomass yield after 2 (Fig. 2a) and 6 weeks (Fig. 2b). The graphs were divided into quadrants based on the growth and pH.

**Fig. 2.**
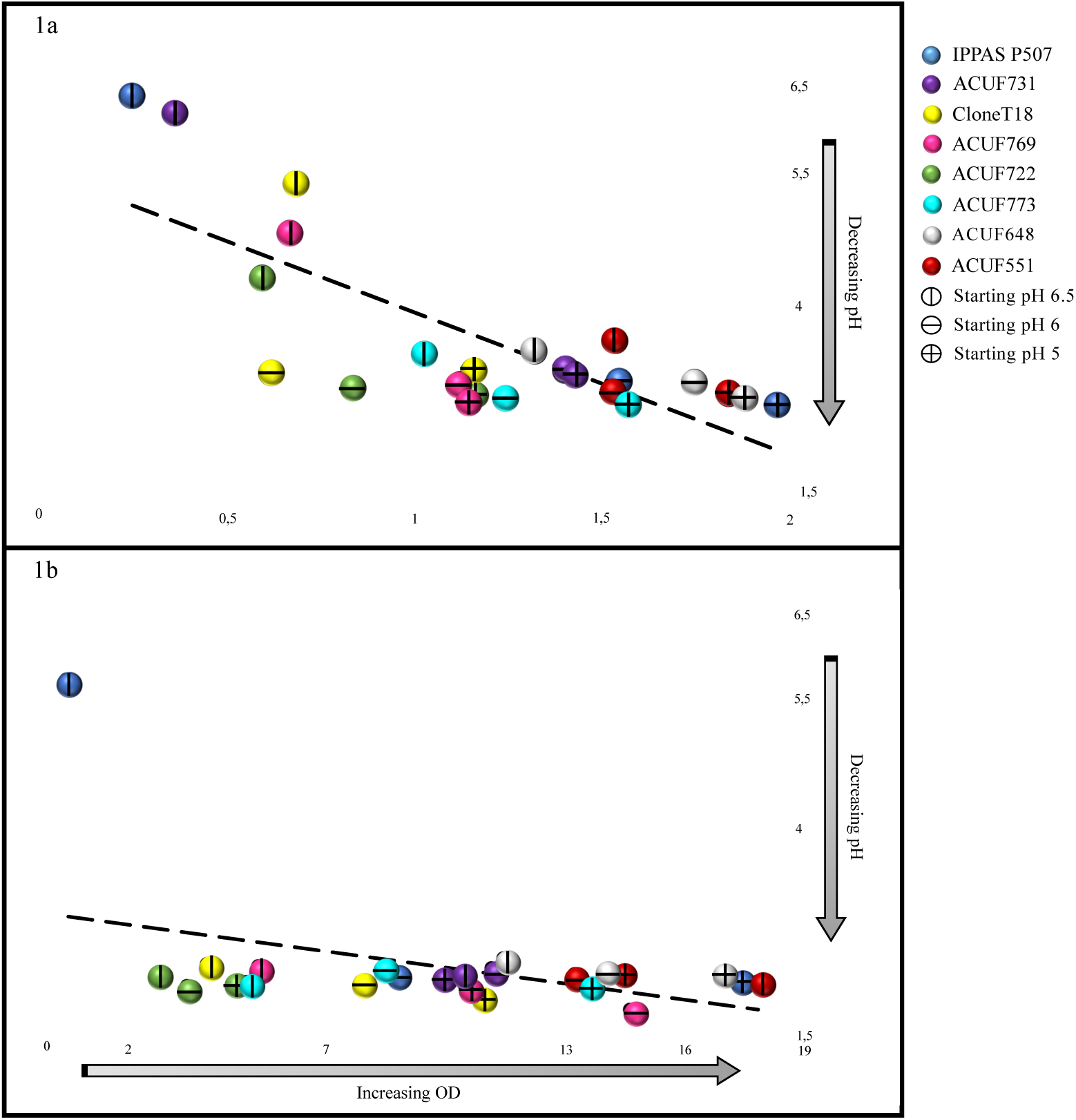
Relationship between pH (y-axis) and growth rate (x-axis) for all the strains after two weeks (1A) and six weeks (1B). correlation = −0.75 (2weeks), −0.43 (6 weeks).

A significant correlation was recorded between pH and growth rate after 2 weeks (corr = - 0.75; P < 0.05). Neither IPPAS P507 nor ACUF731 grew (OD < 0.5) or lowered pH of the medium at starting pH 6.5. Strains ACUF722, ACUF769, and CloneT18 slightly decreased the pH from 6.5 to values ranging from 4 to 5.5, resulting in a moderate increase in biomass (0.5 < OD < 1). Strains ACUF773 and ACUF648 reached the highest biomass yields (1 < OD < 1.5,), as a result of growing acidification of the external medium (pH< 4). Strain ACUF551 decreased the pH to more acidic value, as well, and exhibited the best growth rate, exceeding 1.5 OD after two weeks.

From an initial pH 6, all strains were able to lower the pH to similar values (pH < 4), although with growth rates significantly variable among them. ACUF722 and CloneT18 did not exceed OD 1 after 2 weeks, whereas strains ACUF773, ACUF769, and ACUF731 had an OD ranging from 1 to 1.5. The highest biomass yields were reached by ACUF648, ACUF551 and IPPAS P507, with OD > 1.5.

At pH 5, all *G. maxima* strains exceeded OD 2 and lowered pH below 4. Noteworthy was that strains IPPAS 507, ACUF648, ACUF551, and ACUF773 reached the highest biomass yield (1.5 < OD < 2). At this starting pH level, the media conditions were sufficient for the *G. maxima* to alter conditions to maximal growth capacities.

After six weeks, all strains, regardless of the initial pH, strongly acidified the external medium (final pH < 3), except IPPAS P507, confirming, therefore, its disability to lower the pH (Fig. 2b). However, the differences in growth rate among strains were still more accentuated (0.7 < OD < 18). From an initial pH 6.5, the best growth rate was recorded by ACUF551 (OD = 17.960, Fig. 2b), while ACUF722, CloneT18, ACUF773 and ACUF769 had a slower growth rate (OD = 2.955, 4.21, 5.12, 5.465, respectively; Fig. 2b). From pH 6, ACUF769 and ACUF722 reached the highest (OD = 14.820) and the lowest (OD = 3.695) biomass yields, respectively; the best growth performances at initial pH 5 were recorded by IPPAS P507 (17.640), while ACUF722 again did not exceed OD 5.

### 3.3 Effect of pH and nitrogen source on growth and chlorophyll content

The combined influence of nitrogen source and pH was tested on three *G. maxima* strains displaying different pH responses (Fig. 3). We chose ACUF551, whose final biomass yield was comparable to its control, ACUF722, which showed slow growth in all pH conditions, compared to pH1.5, and IPPAS P507 as the only strain unable to lower the external pH from 6.5.

**Fig. 3.**
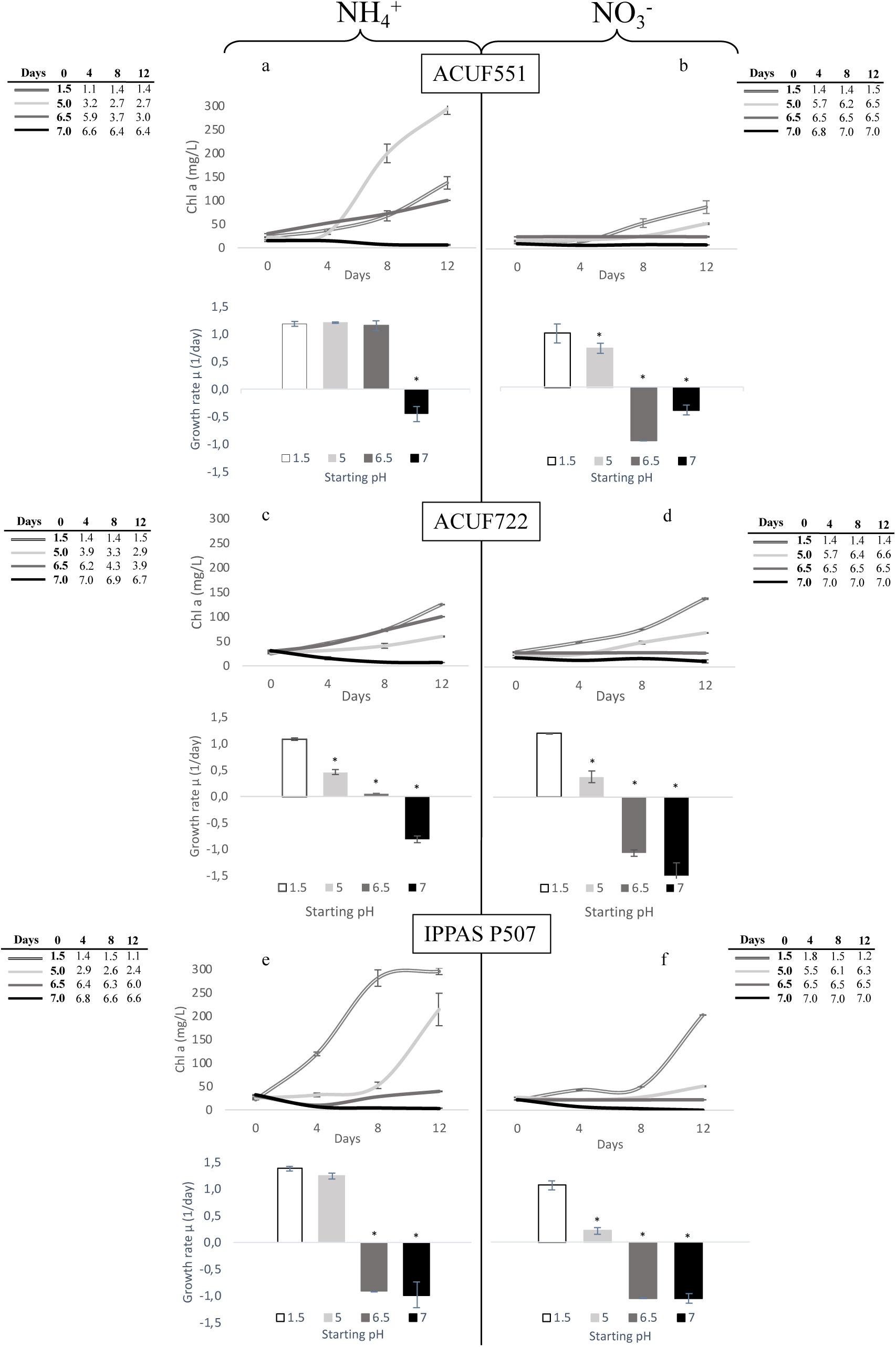
Trend of chlorophyll content during 12 days. The column charts indicate the growth rate measured at the end of the experiments. Data are shown as the mean value (± S.D.), n= 3; Values marked with asterisk are significantly different from the control (Tukey test; p < 0.05).

*G. maxima* ACUF551 grew similarly on either ammonium and nitrate, at pH 1.5, even if the chlorophyll content was significantly higher on NH_4_^+^ than on NO_3_^-^ (Figs. 3a, b; 137.15mg/L on NH_4_^+^ *vs* 76.23mg/L on NO_3_^-^; P < 0.05). After 2 weeks from an initial pH 5 on ammonium, chlorophyll content was 2.14 fold compared to the control (293.33mg/L; P < 0.05); the algal strain did not exhibit difference in growth rates at pH 5 and 6.5, compared to the control; acidification of the medium progressively occurred, reaching 2.7 and 3.0, respectively. Interestingly, *G. maxima* ACUF551 showed the ability to grow on nitrate at pH 5, despite the progressive increasing pH of the medium, which reached 6.5 after two weeks. Chlorophyll levels were detected to increase throughout the experiments, despite achieving values lower than in the controls.

Comparable growth rates and chlorophyll contents were observed in *G. maxima* ACUF722 on both nitrogen sources at pH 1.5 (Figs. 3c, d). A significant decrease in growth rate and chlorophyll content was recorded at pH 5 on ammonium, despite the progressive acidification of the medium (final pH 2.9) as well as on nitrate, where alkalization took pH at 6.6.

As for ACUF551, no significant differences were observed in IPPAS P507 growth rate, when cultivated at pH 1.5 or pH 5 (final pH 2.4) on ammonium (Fig. 3e). From an initial pH 5, chlorophyll levels increased reaching 214mg/L after a lag phase in the first 8 days, progressively acidifying the medium. When the medium was supplemented with nitrate instead of ammonia, the progressive increase of pH from 5 up to 6.3 strongly affected both the growth rate and the chlorophyll content (Fig. 3f).

Negative growth rates were observed for all strains, neither at pH 6.5 on nitrate or at pH 7 on both N-sources.

### 3.4 Effect of pH and nitrogen source on photochemical activity

Photochemical performances were evaluated by rapid and non-invasive PAM fluorometry (Figs. 4-6, Figs. S1-6). From an initial pH 1.5 with either nitrogen sources, the efficiency of photochemical activity was optimal and comparable. Under ammonium as the nitrogen source, ACUF551 had Fv/Fm values around 0.5-0.6; in *G. maxima* strains ACUF722 and IPPAS P507 this variable was stable around 0.6 (Fig. 4a, c, e). On nitrate, all strains resulted in values of Fv/Fm ≥ 0.6 (Fig. 4b, d, f). Maximum quantum yield was recorded at 31 μmol photons m^−2^ s^−1^, both on ammonium and on nitrate (0.4, Fig. 6a-f, Figs. S4-6), with relative low NPQ values (0.05-0.08, Fig. 5a-f, Figs. S1-3); the photochemical activity slightly decreased with increasing light intensities, reaching the minimum value at 281 μmol photons m^−2^ s^−1^ (Fig. 6a-f, Figs. S4-6), where NPQ achieved values ranging from 0.8 to 1 (Fig. 5a-f, Figs. S1-3).

**Fig. 4.**
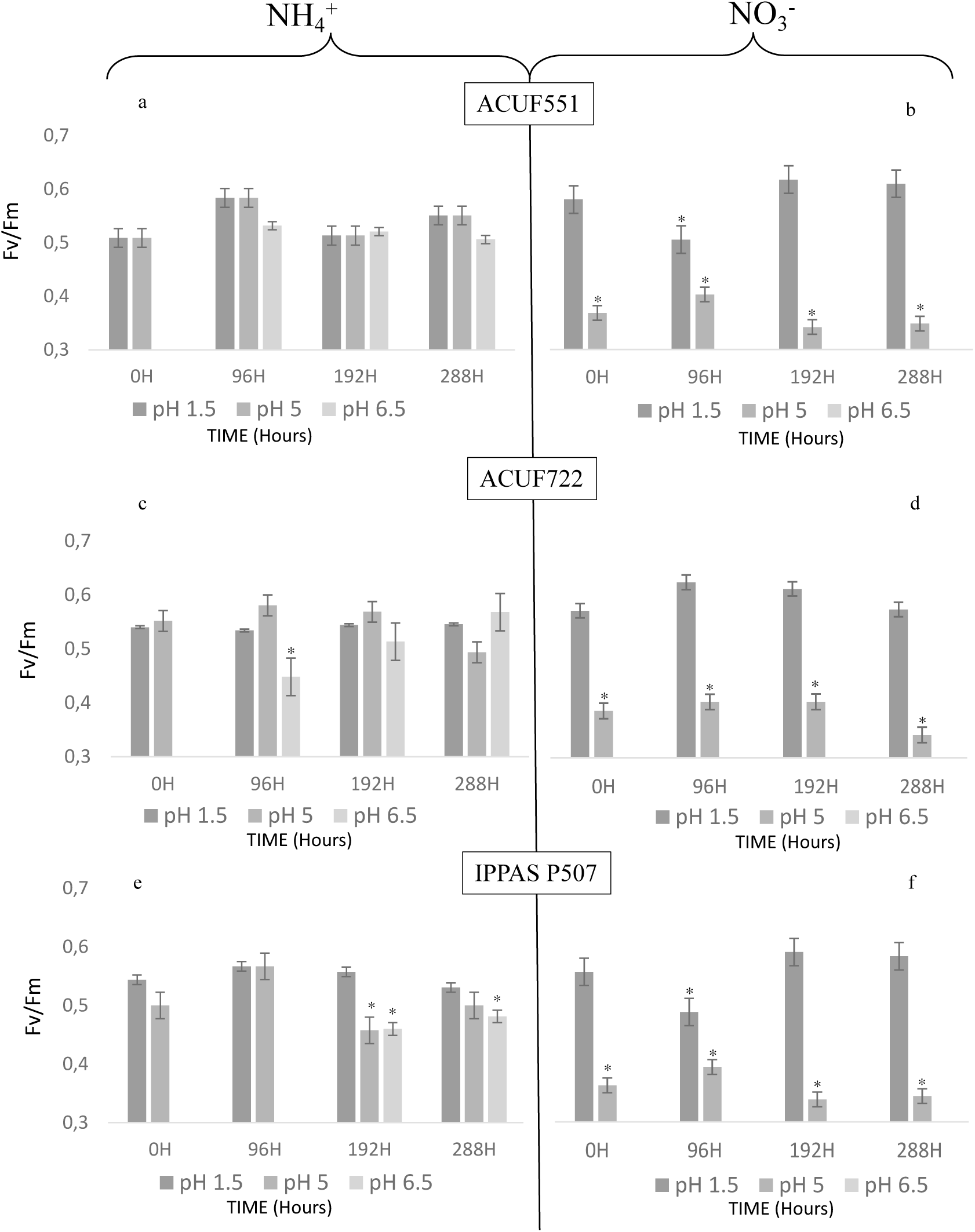
Fv/Fm values of the three G. maxima strains in presence of ammonium and nitrate under different pHs; the quantum efficiency of photosystem II (Fv/Fm) was recorded after 0, 96, 192 and 288h. Data are shown as the mean value (± S.D.), n= 3; p < 0.05.

**Fig. 5.**
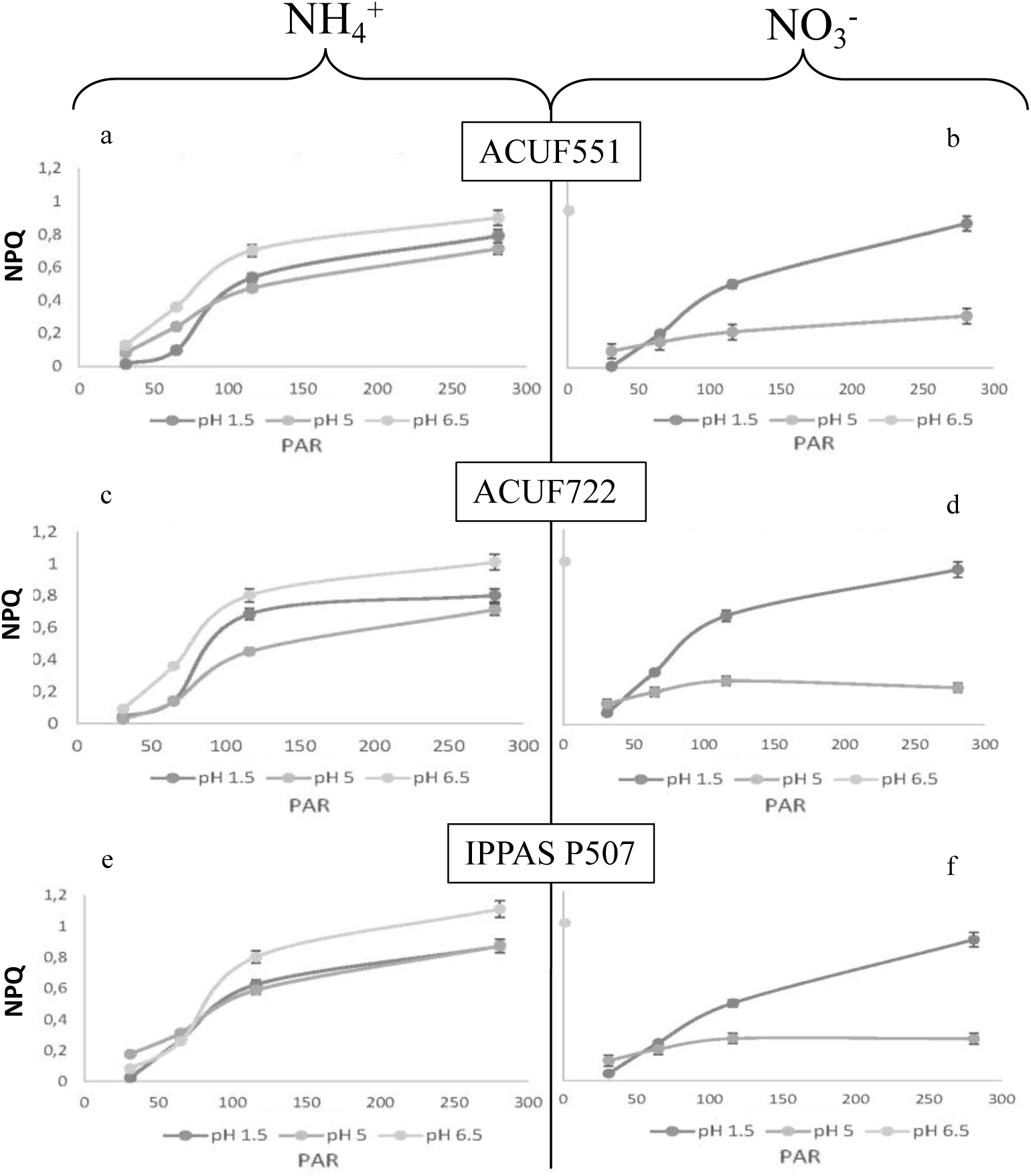
NPQ values measured under ammonium and nitrate source under different pHs, at different light intensities after 288h. Data are shown as the mean value (± S.D.), n= 3; Values marked with asterisk are significantly different from the control (Tukey test; p < 0.05).

**Fig. 6.**
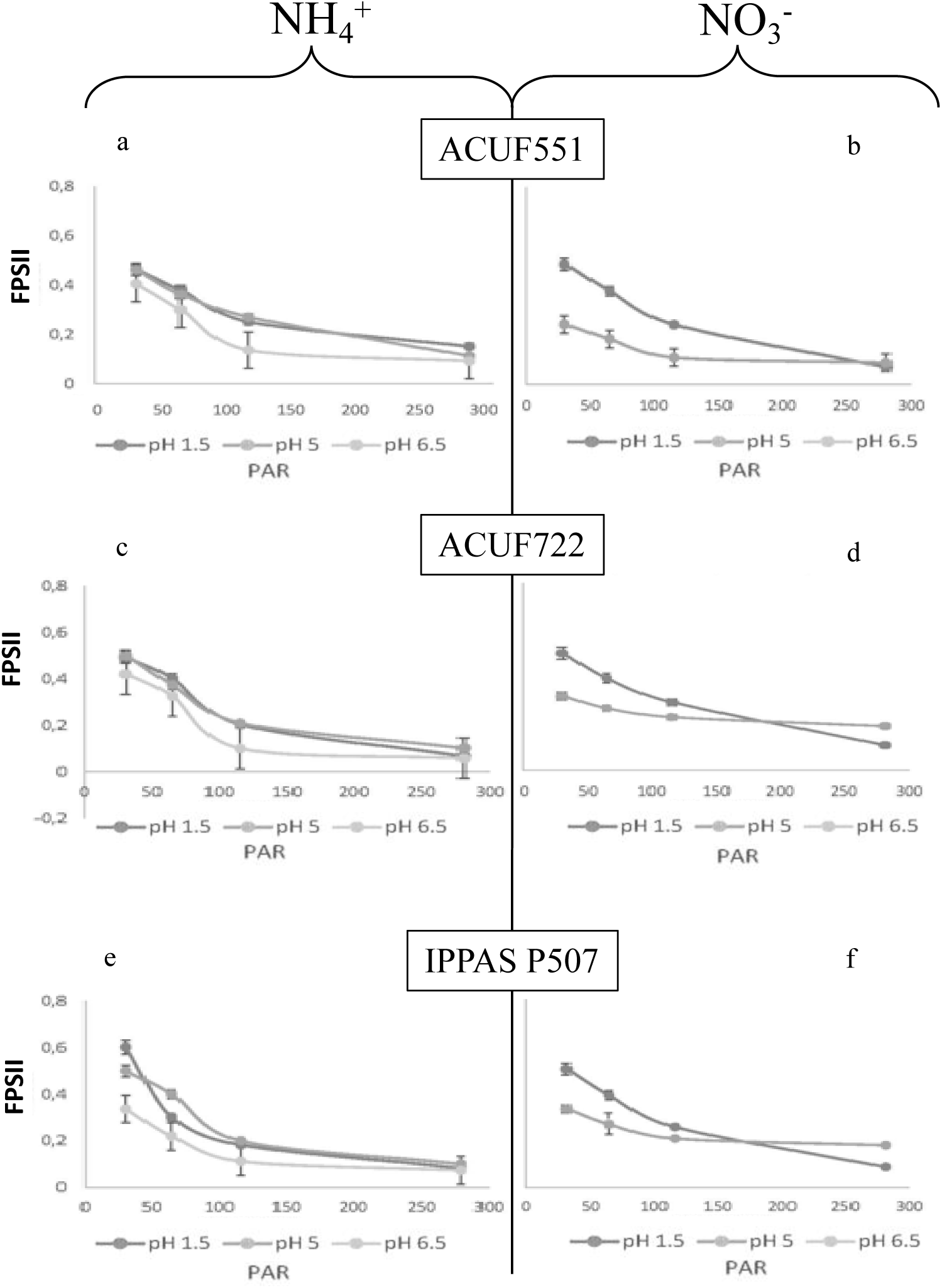
FPSII values measured under ammonium and nitrate source under different pHs, at different light intensities after 288h. Data are shown as the mean value (± S.D.), n= 3; Values marked with asterisk are significantly different from the control (Tukey test; p < 0.05).

Different photochemical performances were noted from a starting pH 5 among the strains on both nitrogen sources: the photochemical efficiency was comparable to pH 1.5, on NH_4_^+^ (Fv/Fm ≥ 0.5, Fig. 4a, c, e), while on nitrate Fv/Fm was significantly lower than ammonium sources (Fv/Fm ≤ 0.4; p < 0.05; Fig. 4b, d, f). Only small variations of NPQ and FPSII were recorded even at the high light intensity of 281 μmol photons m^−2^s^−1^ (Figs. 5 and 6 a-f, Figs. S1-6).

From pH 6.5, the photochemical activity of ACUF551 and ACUF722 on ammonium was inhibited until 96 hours. After this time as an acclimation period, physiological activity started again (Fig. 4a, c). The Russian IPPAS P507 showed an acclimation period of 192 hours on ammonium, showing a lower photochemical activity compared to the other strains (Fig. 4e). No photochemical activity was recorded on nitrate at pH 6.5 in all strains, and neither at pH 7 in all the isolates in the presence of both nitrogen sources.

## DISCUSSION

We applied different methods to assess the influence of pH and nitrogen source on the physiology of eight *G. maxima* strains. Our results showed a significant correlation between the growth rates and the pH lowering on NH_4_^+^medium in different strains collected from acidic and non-acidic geothermal environments. It is known that the assimilation of ammonium produces protons which must be promptly extruded to avoid cell acidification and support metabolism [37]. All strains used in this work showed comparable abilities in lowering the pH when cultured at pH 5 and 6 in nitrogen provided as NH_4_^+^, although with variable growth rate.

Substantial differences among them were recorded at pH 6.5; the Russian IPPAS P507 required a more extended phase of adaptation, and this strain scarcely grew during the six weeks of examination. A decrease in the external pH reached only the value of 5.7. In contrast, the Icelandic ACUF551 and the Turkish ACUF769, both isolated from acidic sites, showed the best growth performances at pH 6.5 and 6, respectively, during the whole experiment. As reported by Lowell and Castenholz (2013) a different ability in lowering pH was also shown by *C. merolae:* eight strains isolated from Yellowstone were able to reduce the pH of the medium, while none of the three Japanese isolates was competent; the authors also highlighted variations in the final yields, ascribing them both to small differences in inoculum density and in real genetic differences among the competent strains [23]. Among our isolates, two of the Turkish strains (ACUF773, ACUF648) from non-acidic sites quickly lowered the pH to values 3.39-3.42, thus allowing a faster growth, which could be a reliable survival strategy in otherwise hostile environments. According to Rysenbach et al. (2002), in some hot springs, reduced constituents such as hydrogen sulfide, ammonia, and methane are far from equilibrium with their oxidized counterparts such as sulfate, nitrate, and carbon dioxide [38]. Ammonia concentrations strongly vary in these environments, as well as ambient pH. Such ecological variations can produce radically different habitats at a small scale and change rapidly over time as well, selecting for organisms that respond quickly to these geochemical fluctuations.

Another interesting aspect emerging from our experiments is related to the growth in the nitrate-rich medium. All *G. maxima* strains were able to use nitrate as well as ammonium. The dominant nitrogen source in acidic hot springs is ammonium, while nitrate and nitrite are usually generated by the oxidising activity of the microbial vitality [38]. As a consequence, nitrate assimilation is rarely essential in these environments [39]. The enzymes usually involved in nitrate assimilation and conversion are nitrate reductase (NR, nitrate to nitrite), and nitrite reductase (NiR, nitrite to ammonium). Among Cyanidiophyceae, *C. merolae* can grow on both nitrogen sources [39]. However, no typical NiR coding gene was found in its complete genome [40]. According to Imamura et al. (2010), the activity is encoded by two candidate genes for sulfite reductase (SiR), located next to the nitrate-related genes. These transcripts are inhibited by ammonium [39]. Similarly, the species *G. sulphuraria* lacks the common NR-coding gene, thus suggesting again the existence of an unusual assimilatory system (Weber unpublished sited in Imamura et al. 2010). By phylogenetic analysis, *G. maxima* is more strictly related to *C. merolae* than to *G. sulphuraria*, despite the striking morphological divergences among the two genera [9,12,13]. It is intriguing that *G*. *maxima* is provided by a nitrate assimilation gene toolkit similar to that of *C. merolae*, thus supporting their phylogenetic relationships (H.S. Yoon, personal communication).

In our pH 1.5 experiments, all cultures were able to use nitrate in a manner significantly comparable to ammonium. These maintained a relatively constant external pH. Photosynthetic efficiency was productive from a low light intensity recorded in all *G. maxima* cultures at pH 1.5 for either in nitrate or ammonium, showing Fv/Fm values around 0.500. The maximum efficiency of photosystem II in the dark (Fv/Fm) usually provides indications on the physiological state of the cultures. Red algae exhibit an optimum Fv/Fm that is lower than the ones manifested by green and brown microalgae (0.700). This reduction is related to the presence of phycobiliproteins [11] in the light harvesting system coupled to PSII, interfering with fluorescence signal [41]. A steady rise of the pH is the result of nitrate uptake and successive transformation to ammonium, requiring a net influx of protons from the culture medium [37]. Cultivation on nitrate from an initial pH 5, and the subsequent increase of the medium pH, correlated with the photochemical activity. Lower values of Fv/Fm, NPQ and FPSII, suggested a stress condition that could influence the protein expression and enzymatic concentration of photosynthetic apparatus [42].

Fluorometric analyses were used to demonstrate an exponential increase in the growth rate at the beginning of our experiments (0-4 days). These remained constant from the 4th day onward; the survival of the cultures is potentially ascribable to switching from the autotrophic metabolism to a heterotrophic one. Oesterhelt et al. (2007) assessed that autotrophic cultures of *G. sulphuraria* (strain 074) exponentially grew up to pH 5 and reduced the growth significantly at higher pHs. Diversely, heterotrophic cultures of the same strain exhibited cell growth up to pH 8 [43].

Environmental pH is one of the most critical factors limiting the dispersal of extremophiles and more specific acidophiles [44]. The discovery of *G. maxima* strains from non-acidic thermal areas and the capacity to tolerate a wide range of initial environmental pHs are strong evidence of their potential adaptation to changing environments. A wide pH range tolerance is a critical ecological factor in the regulation of the nutrient cycle (*i.e.* carbon, nitrogen, and phosphorus) under extreme environmental conditions. Several studies already reported the ability of microorganisms collected from acidic or alkaline environments to survive in a wide range of pH, suggesting these capacities as indicators of the ecological resilience of extremophiles [44]. According to Dhakar and Pandey (2016), survival in sudden adverse environmental conditions is the consequence of the expression of “hidden” genomic information (Cryptic Genetic Variation, CGV). The changes in ecological factors induce the expression of hidden genes without any alteration in the genetic composition or proper effect on phenotype; transcriptomics and proteomics can be helpful to reveal the change in the cellular response to pH changes [1]. The ability of the extremophiles to respond at environmental variations, such as pH, lies in the expression and production of proteins, extremozymes or secondary metabolites. These compounds are structurally suitable and of a broad interest in biotechnological processes, including both environmental (bioremediation, biodegradation, and biocontrol), and industrial applications. We see these as sources of new extremozymes and other useful bioactive compounds with wide pH range tolerance.

## ACKNOWLEDGEMENTS

Special thanks to the Algal Collection University Federico II, ACUF (www.acuf.net) for the contribution in the maintenance of the algal isolates.

## DECLARATION OF AUTHOR CONTRIBUTIONS

M. Iovinella contributed to the conception, analysis, interpretation of data and drafting; D.A. Carbone was involved in the photochemical investigations; D. Cioppa contributed to the experimental execution; M. Innangi was involved in the statistical analysis and interpretation of data; S. Esposito provided resources (instrumentation, reagents and analysis tools); S.J. Davis and C. Ciniglia brought a critical revision of the article and approved the submission.

