## Supplementary Figures for "Prevalent pH Controls the Capacity of *Galdieria Maxima* to Use Ammonia and Nitrate as a Nitrogen Source"

**Fig.S1** NPQ values measured under ammonium and nitrate source at different light intensities at the beginning of the experiment

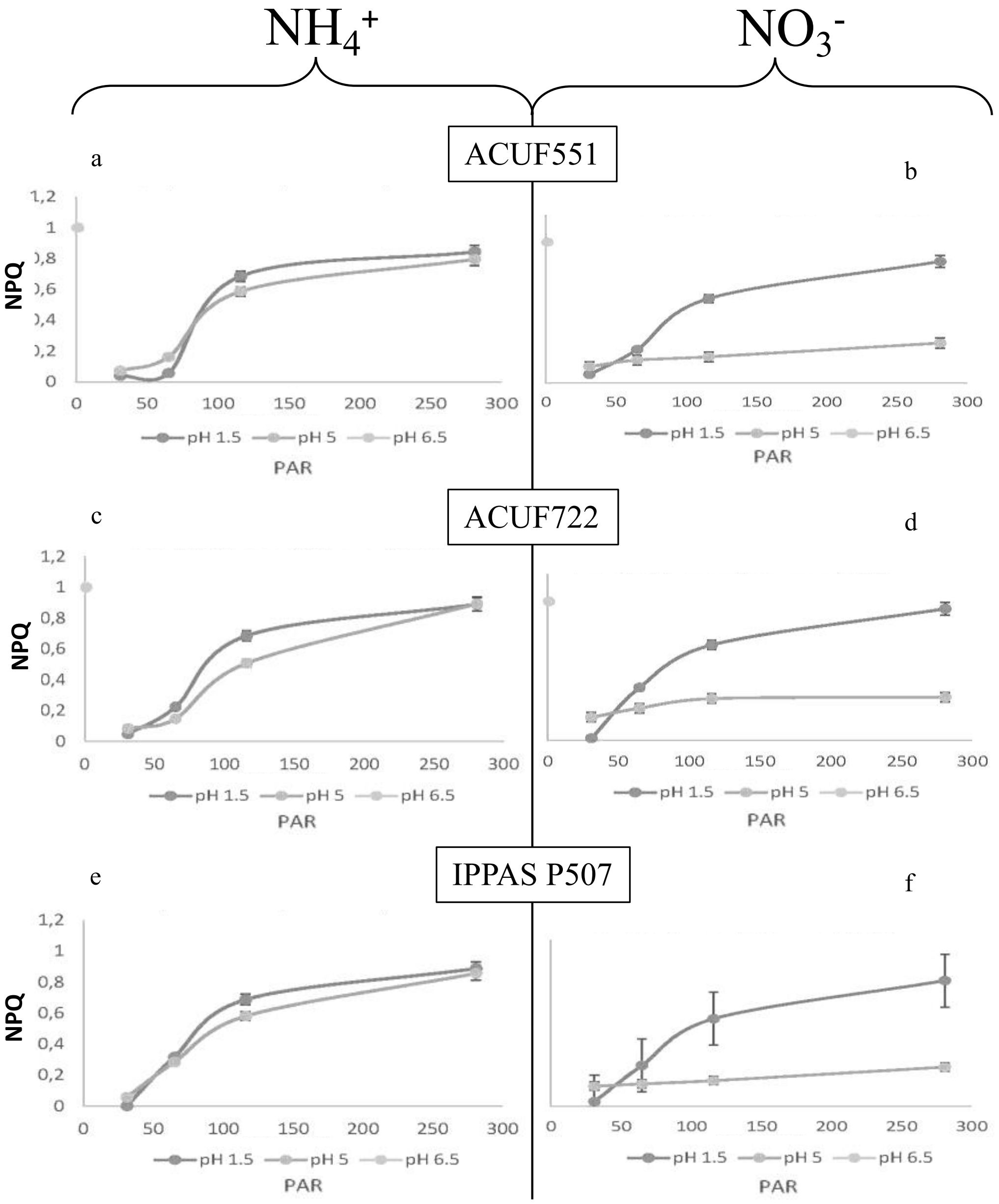

**Fig.S2** NPQ values measured under ammonium and nitrate source at different light intensities after 4 days

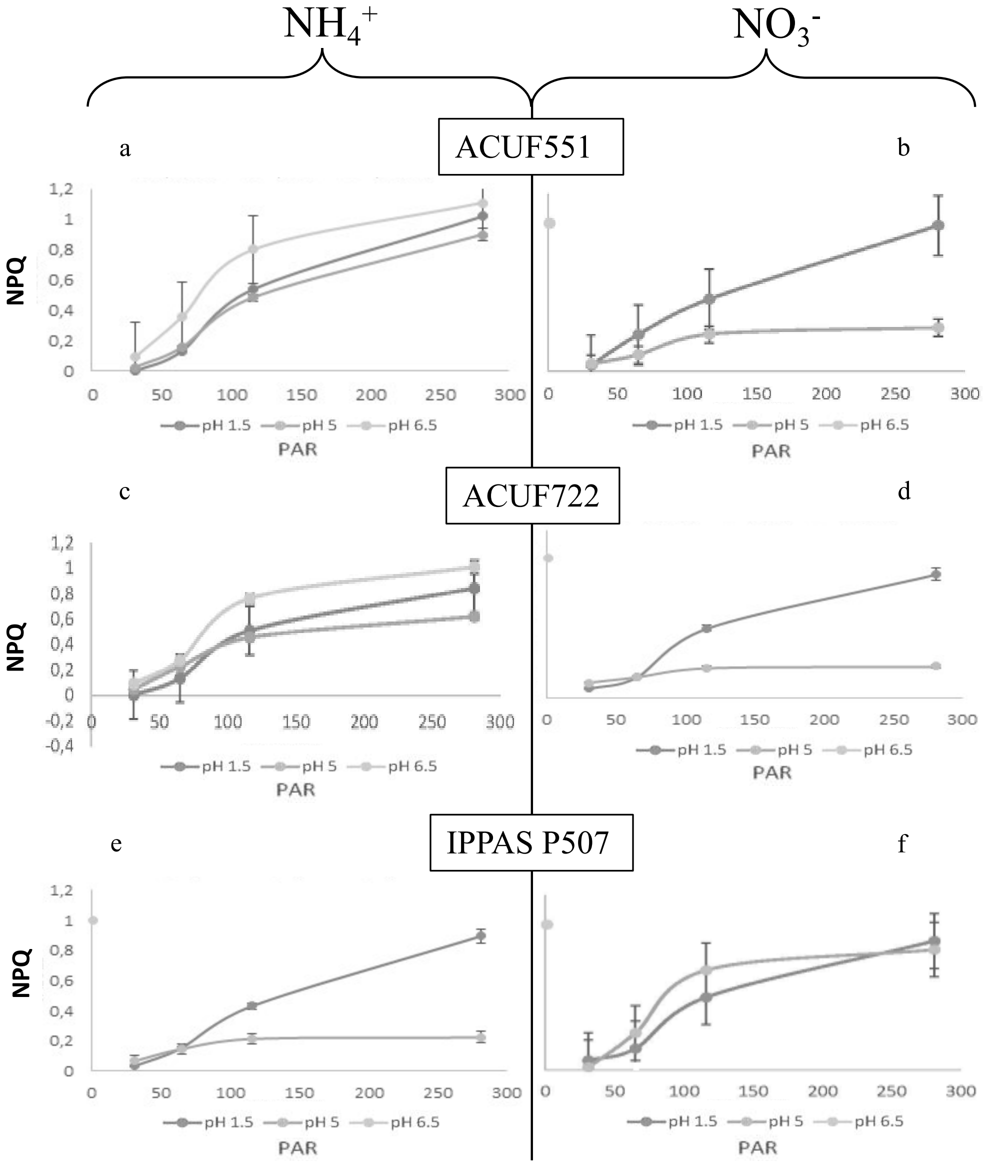

**Fig.S3** NPQ values measured under ammonium and nitrate source at different light intensities after 8 days

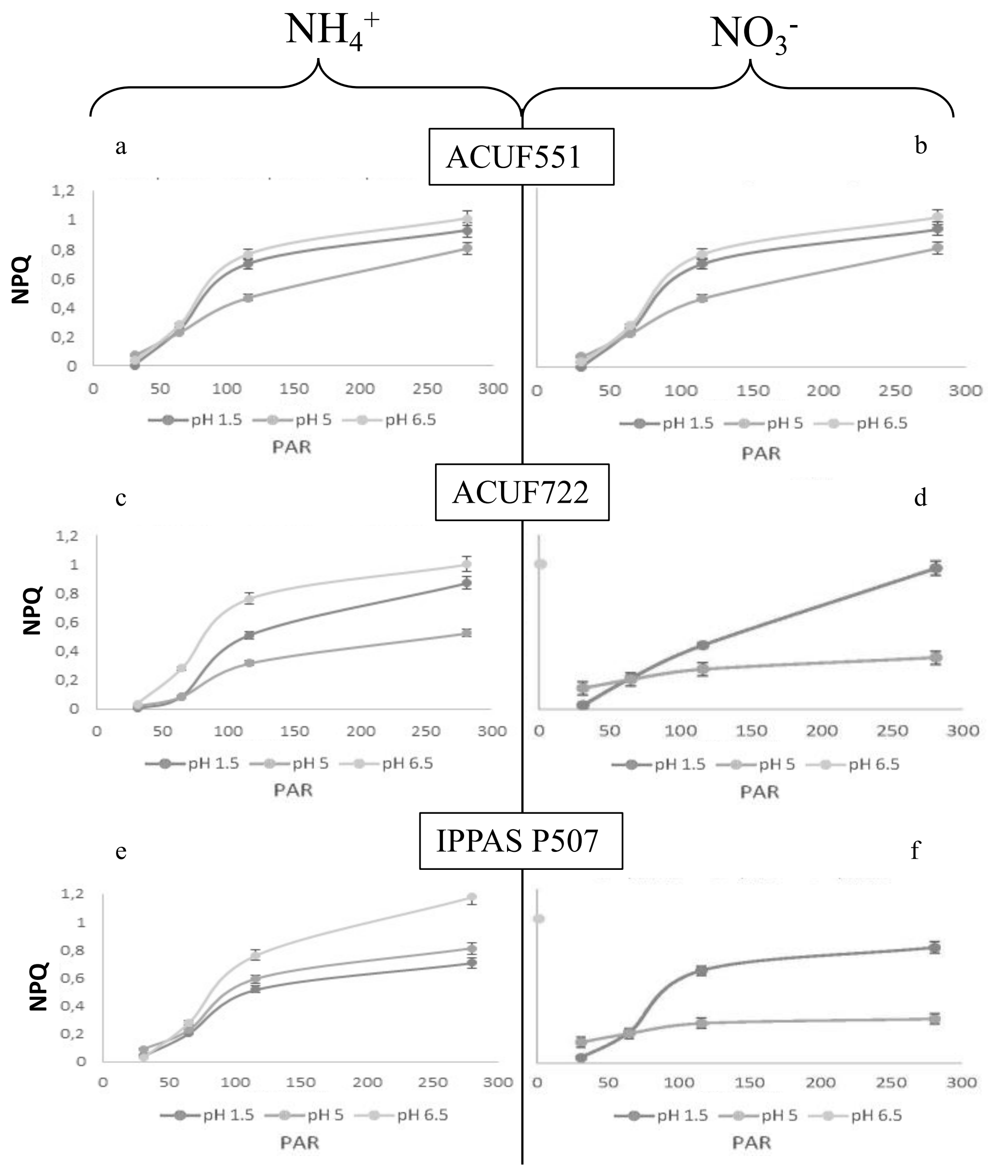

**Fig.S4** FPSII values measured under ammonium and nitrate source at different light intensities at the beginning of the experiment

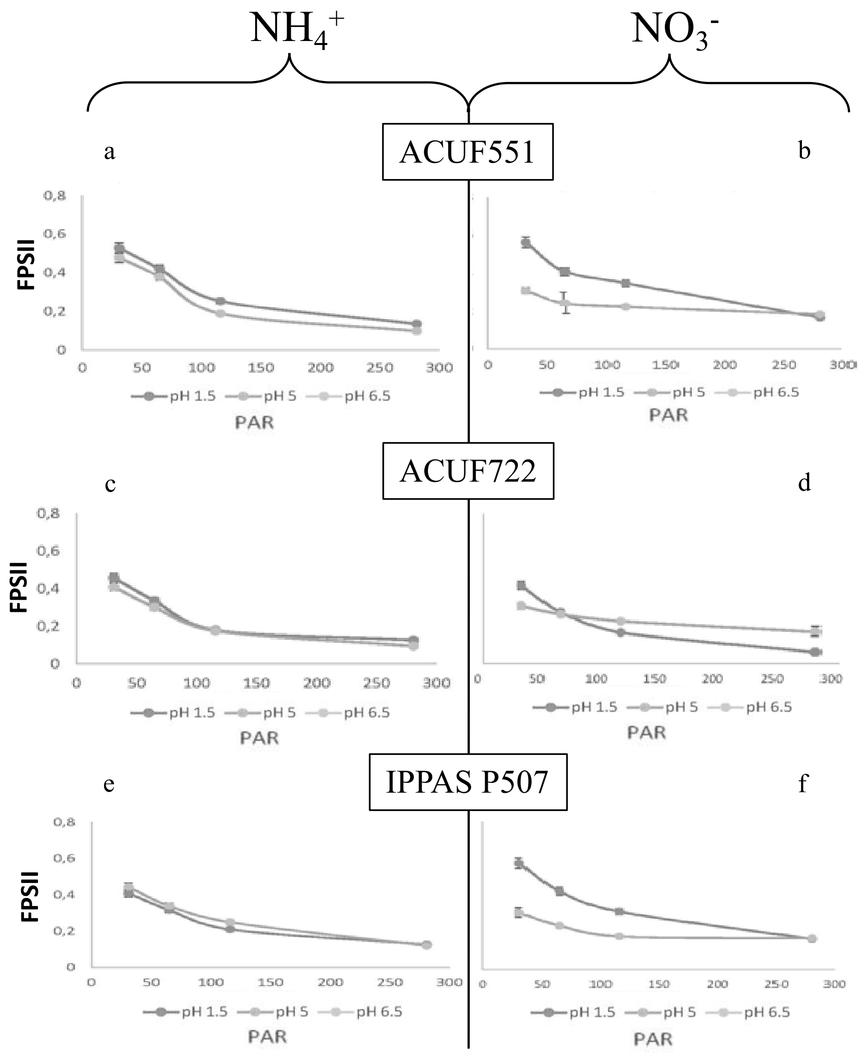

**Fig.S5** FPSII values measured under ammonium and nitrate source at different light intensities after 4 days

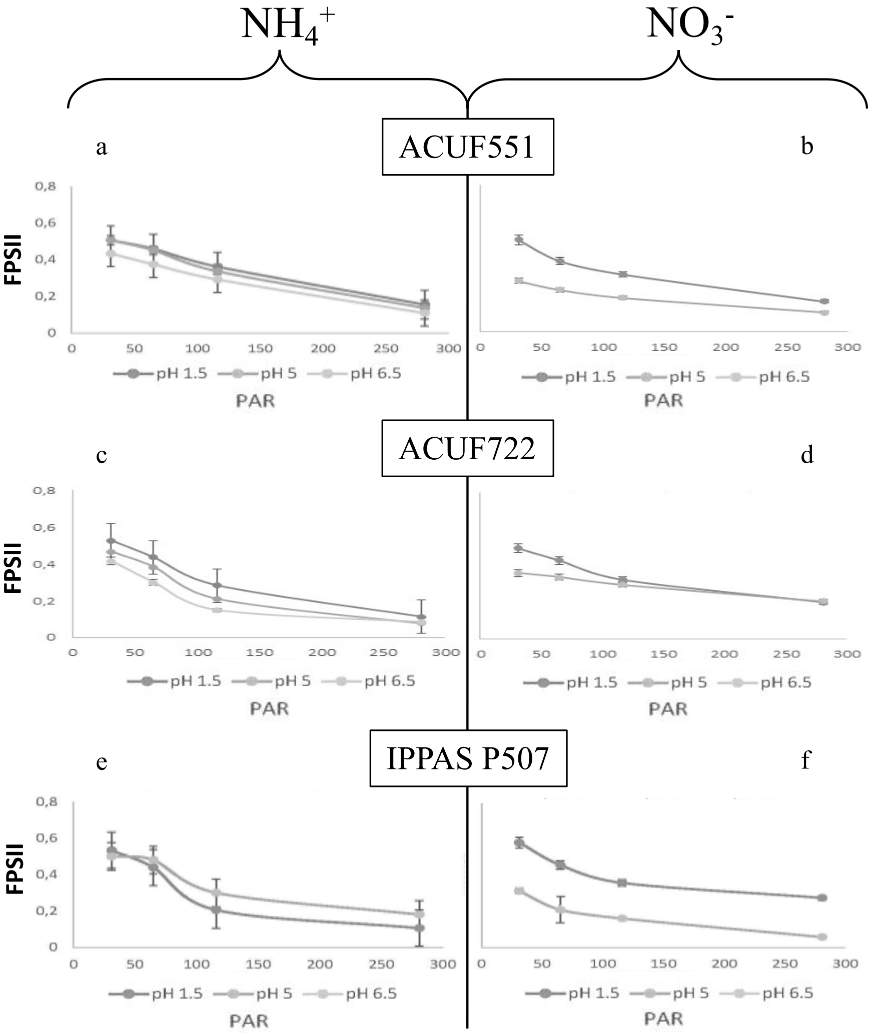

**Fig.S6** FPSII values measured under ammonium and nitrate source at different light intensities after 8 days

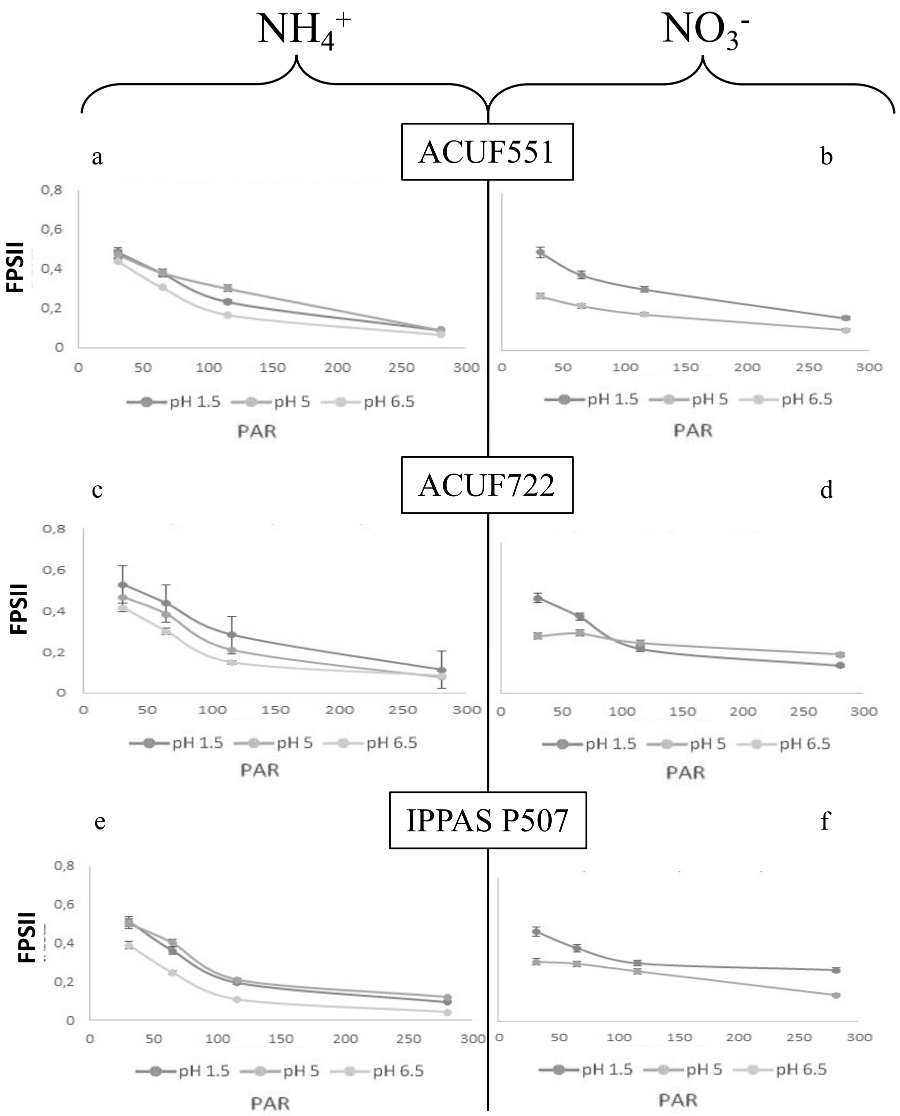
