## Supplementary Tables for "Prevalent pH Controls the Capacity of *Galdieria Maxima* to Use Ammonia and Nitrate as a Nitrogen Source"

**Tab.S1** Taxa, collection site, strain voucher and accession numbers (rbcL) for the dataset.

| Taxa | Collection site | Strain Voucher | Accession Number |
|---|---|---|---|
| *Galdieria maxima* | Kunashir (Russia) | IPPAS P507 | AY391370 |
| | Landmannalaugar (Iceland) | ACUF551 | MK394995 |
| | | ACUF419 | KC883840 |
| | | ACUF420 | KC883841 |
| | | ACUF421 | KC883842 |
| | | ACUF428 | KC883848 |
| | | ACUF449 | KC883861 |
| | | ACUF450 | KC883862 |
| | | ACUF456 | KC883868 |
| | | ACUF457 | KC883869 |
| | | ACUF458 | KC883870 |
| | | ACUF451 | KC883863 |
| | Seltun (Iceland) | ACUF389 | KC883816 |
| | | ACUF393 | KC883819 |
| | | ACUF396 | KC883821 |
| | | ACUF425 | KC883846 |
| | | ACUF436 | KC883849 |
| | | ACUF468 | KC883880 |
| | | ACUF469 | KC883881 |
| | | ACUF411 | KC883833 |
| | Niasjvellir (Iceland) | ACUF404 | KC883827 |
| | Güçlükonak (Turkey) | ACUF769 | KX501179 |
| | | ACUF722 | KX501174 |
| | | ACUF660 | KY033404 |
| | | ACUF710 | KX501173 |
| | | ACUF697 | KY033415 |
| | Diyadin (Turkey) | cloneT18 | KX501185 |
| | | ACUF773 | KX501180 |
| | | ACUF665 | KY033406 |
| | | ACUF774 | KY033436 |
| | | cloneT06 | KY033453 |
| | | ACUF772 | KY033435 |
| | Manisa Kula (Turkey) | ACUF648 | KY033397 |
| | | ACUF731 | KY033420 |
| | | ACUF671 | KY033410 |
| | | ACUF743 | KY033428 |
| | | ACUF777 | KY033439 |
| | | ACUF776 | KY033438 |
| | Germencik (Turkey) | cloneT15 | KX501183 |
| | | ACUF673 | KY033411 |

| | | | |
|---|---|---|---|
| | | ACUF739 | KY033425 |
| | | ACUF742 | KY033427 |
| | Cermick (Turkey) | ACUF647 | KY033396 |
| | | cloneT03 | KY033450 |
| | | cloneT04 | KY033451 |
| | | ACUF650 | KY033398 |
| | | ACUF766 | KY033431 |
| | | ACUF783 | KY033443 |
| | Biloris (Turkey) | ACUF653 | KY033400 |
| | | ACUF763 | KY033429 |
| | | ACUF764 | KY033430 |
| | | ACUF735 | KY033422 |
| *Galdieria sulphuraria* | Yellostone National Park (USA) | SAG108.79 | AY119767 |
| | California (USA) | UTEX2393 | AF233069 |
| | Caserta (ITaly) | ACUF011 | AY541303 |
| | Los Azufres (Mexico) | ACUF135 | AY541309 |
| | Benevento (Italy) | ACUF012 | AY541310 |
| | Solfatara (Italy) | ACUF017 | AY541306 |
| | Scarfoglio (Italy) | ACUF018 | AY541304 |
| | Vulcano (Italy) | ACUF021 | AY541307 |
| | Ischia (Italy) | ACUF015 | AY541305 |
| | Sasso Pisano (Italy) | SP3-C2 | DQ916749 |
| | Sasso Pisano (Italy) | SP1-10 | DQ916748 |
| | Monte Rotondo (Italy) | MR6-C36 | DQ916747 |
| | | MR5-C17 | DQ916746 |
| | | MR4-21 | DQ916745 |
| | Pisciarelli (Italy) | cloneA12 | AY541313 |
| | | cloneD5 | AY541321 |
| | | cloneD15 | AY541322 |
| | | cloneE11 | AY541324 |
| | | cloneE12 | AY541325 |
| | Gunnhuver (Iceland) | ACUF381 | KC883808 |
| | | ACUF382 | KC883809 |
| | Landmannalaugar (Iceland) | ACUF385 | KC883812 |
| | | ACUF386 | KC883813 |
| | | ACUF387 | KC883814 |
| | | ACUF388 | KC883815 |
| | Seltun (Iceland) | ACUF395 | KC883820 |
| | | ACUF397 | KC883822 |
| | | ACUF398 | KC883973 |
| | | ACUF422 | KC883843 |
| | | ACUF423 | KC883844 |
| | | ACUF424 | KC883845 |

| | | | |
|---|---|---|---|
| | | ACUF437 | KC883850 |
| | | ACUF409 | KC883831 |
| | | ACUF448 | KC883860 |
| | | ACUF452 | KC883864 |
| | | ACUF439 | KC883852 |
| | | ACUF454 | KC883866 |
| | | ACUF440 | KC883853 |
| | | ACUF470 | KC883882 |
| | | ACUF472 | KC883883 |
| | | ACUF473 | KC883884 |
| | | ACUF474 | KC883885 |
| | | ACUF475 | KC883886 |
| | | ACUF459 | KC883871 |
| | | ACUF460 | KC883872 |
| | | ACUF410 | KC883832 |
| | | ACUF463 | KC883875 |
| | | ACUF417 | KC883839 |
| | | ACUF412 | KC883834 |
| | Niasjvellir (Iceland) | ACUF399 | KC883823 |
| | | ACUF400 | KC883824 |
| | | ACUF443 | KC883855 |
| | | ACUF444 | KC883856 |
| | Viti (Iceland) | ACUF461 | KC883873 |
| | Güçlükonak (Turkey) | ACUF658 | KY033403 |
| | | ACUF725 | KX501175 |
| | | ACUF768 | KX501178 |
| | | ACUF781 | KY033441 |
| | Germencik (Turkey) | ACUF778 | KX501181 |
| | | ACUF674 | KY033412 |
| | | ACUF676 | KY033413 |
| | | ACUF779 | KY033440 |
| | | cloneT02 | KY033449 |
| *Galdieria partita* | Kamchatka (Russia) | IPPAS P500 | AB18008 |
| *Galdieria daedala* | Kunashir (Russia) | IPPAS P508 | AY541302 |
| *Galdieria phlegrea* | Viterbo (Italy) | ACUF009 | AY119768 |
| | Agrigento (Italy) | ACUF063 | AY119769 |
| | Pisciarelli (Italy) | ACUF002 | AY541311 |
| | | cloneB15 | AY541314 |
| | | cloneB19 | AY541315 |
| | | cloneB20 | AY541316 |
| | | cloneC1 | AY541317 |

| | | | |
|---|---|---|---|
| | Biloris (Turkey) | ACUF765 | KX501177 |
| | | ACUF652 | KY033399 |
| | | ACUF657 | KY033402 |
| | | ACUF780 | KX501182 |
| | | ACUF656 | KY033401 |
| | Güçlükonak (Turkey) | ACUF784 | KY033444 |
| | | cloneT09 | KY033456 |
| | | cloneT16 | KX501184 |
| | Nemrut (Turkey) | ACUF738 | KX501176 |
| | | ACUF664 | KY033405 |
| | Diyadin (Turkey) | ACUF667 | KY033407 |
| | | ACUF669 | KY033409 |
| | | ACUF788 | KY033447 |
| | | ACUF771 | KY033434 |
| | Cermik (Turkey) | ACUF642 | KY033395 |
| | | ACUF625 | KY033394 |
| | | cloneT07 | KY033454 |
| | | cloneT08 | KY033455 |
| | | cloneT10 | KY033457 |
| | | ACUF668 | KY033408 |
| *Cyanidioschyzon merolae* | Java (Indonesia) | ACUF201 | AY119765 |
| | Monte Nuovo (Italy) | ACUF202 | AY541296 |
| | Pisciarelli (Italy) | ACUF001 | AY119766 |
| | | cloneA1 | AY541312 |
| | | cloneD1 | AY541320 |
| | | cloneE10 | AY541323 |
| | Biloris (Turkey) | cloneT01 | KY033448 |
| | Nemrut (Turkey) | cloneT05 | KY033452 |
| *Cyanidium caldarium* | Siena (Italy) | ACUF019 | AY541297 |
| | Java (Indonesia) | ACUF182 | AY541298 |
| | Acqua Santa (Italy) | ACUF020 | AY541299 |
| | Monte Rotondo (Italy) | MR4-22 | DQ916750 |
| | | MR5-5 | DQ916751 |
| | | MR6-C35 | DQ916752 |
| | Sasso Pisano (Italy) | SP1-10 | DQ916753 |
| | Pisciarelli (Italy) | cloneC2 | AY541318 |
| | Cermik (Turkey) | ACUF767 | KY033432 |
| | Diyadin (Turkey) | ACUF775 | KY033437 |
| | Güçlükonak (Turkey) | cloneT17 | KY033462 |
| *Cyanidium chilense* | Monte Rotaro (Italy) | sp.19 | AY541300 |
| | | sp.20 | AY541301 |
| | Terme di baia (Italy) | sp.21 | KC914876 |
| Cyanidales sp. | Yellowstone National Park (USA) | DS1-9 | JQ269631 |

| | | | |
|---|---|---|---|
| | | DS2-5 | JQ269629 |
| | | DS3-1 | JQ269633 |
| | | SFFL-5 | JQ269630 |
| | | LCATERR-7 | JQ269608 |
| | | LCBCEL-5 | JQ269609 |
| | | LCBTERR-6 | JQ269612 |
| | | CHJ-4 | JQ269635 |
| | | CHJ-5 | JQ269634 |
| | | DSB-9 | JQ269617 |
| | | DSC-8 | JQ269618 |
| | | DSD-7 | JQ269605 |
| | | DSE-8 | JQ269619 |
| | | DSF-12 | JQ269620 |
| | | DSH-4 | JQ269606 |
| | | DS1-6 | JQ269623 |
| | | DS2-2 | JQ269638 |
| | | DS3-3 | JQ269624 |
| | | SFFL-8 | JQ269630 |
| | | SFFR-7 | JQ269616 |
| | | LCASUB-11 | JQ269627 |
| | | LCBCEL-7 | JQ269610 |
| | | LCBSUB-5 | JQ269628 |
| | | LCBTERR-12 | JQ269611 |
| | | LCCBLGR-8 | JQ269613 |
| | | LCCYEGR-4 | JQ269614 |
| | | NCB-4 | JQ269621 |
| | | RIVER1B-5 | JQ269636 |
| | | SSI-6 | JQ269622 |
| | | SSII-1 | JQ269626 |
| | | TS-4 | JQ269637 |
| **Outgroups** | | | |
| *Rhodella violacea* | / | SAG115.79 | AY119776 |
| *Bangiopsis subsimplex* | / | PR21 | AY119772 |
| *Dixoniella grisea* | / | SAG39.94 | AY119773 |
| *Porphyridium aerugineum* | / | SAG1380-2 | AY119775 |

**Tab.S2** Optical densities and pHs measured weekly during six weeks of growth under ammonium as the nitrogen source starting from pH 7, 6.5, 6, 5 and 1.5.

| **Start pH 7** | | **Time 0** | **Week 1** | **Week 2** | **Week 3** | **Week 4** | **Week 5** | **Week 6** |
|---|---|---|---|---|---|---|---|---|
| IPPAS P507 | OD | 0.343 ± 0.006 | 0,.364 ± 0.001 | 0.353 ± 0.001 | 0.312 ± 0.002 | 0.313± 0.005 | 0.317± 0.001 | 0.372 ± 0.004 |
| | pH | 7.00 ± 0.02 | 7.01 ± 1.10 | 6.98 ± 0.90 | 7.02 ± 0.60 | 6.97 ± 0.70 | 6.99 ± 0.60 | 7.01 ± 0.50 |
| ACUF769 | OD | 0.335 ± 0.006 | 0.277 ± 0.002 | 0.317 ± 0.001 | 0.313 ± 0.005 | 0.360 ± 0.018 | 0.313 ± 0.005 | 0.310 ± 0.001 |
| | pH | 6.46 ± 0.01 | 6.05 ± 0.11 | 4.76 ± 0.62 | 3.16 ± 0.11 | 2.80 ± 0.08 | 2.52 ± 0.05 | 2.33 ± 0.03 |
| ACUF722 | OD | 0.310 ± 0.001 | 0.257 ± 0.054 | 0.313 ± 0.004 | 0.321 ± 0.005 | 0.326 ± 0.002 | 0.325 ± 0.001 | 0.324 ± 0.011 |
| | pH | 6.50 ± 0.04 | 6.12 ± 0.04 | 4.26 ± 0.13 | 3.05 ± 0.04 | 2.74 ± 0.02 | 2.47 ± 0.01 | 2.25 ± 0.00 |
| CloneT18 | OD | 0.326 ± 0.002 | 0.296 ± 0.029 | 0.324 ± 0.001 | 0.355 ± 0.011 | 0.348 ± 0.001 | 0.335 ± 0.006 | 0.332 ± 0.001 |
| | pH | 6.51 ± 0.09 | 6.29 ± 0.00 | 5.33 ± 0.32 | 3.34 ± 0.16 | 2.81 ± 0.08 | 2.55 ± 0.08 | 2.37 ± 0.06 |
| ACUF773 | OD | 0.335 ± 0.006 | 0.340 ± 0.001 | 0.306 ± 0.005 | 0.343 ± 0.006 | 0.303 ± 0.001 | 0.343 ± 0.008 | 0.346 ± 0.001 |
| | pH | 6.55 ± 0.01 | 5.86 ± 0.04 | 3.39 ± 0.45 | 2.90 ± 0.06 | 2.66 ± 0.03 | 2.44 ± 0.02 | 2.17 ± 0.01 |
| ACUF648 | OD | 0.356 ± 0.001 | 0.351 ± 0.005 | 0.352 ± 0.002 | 0.292 ± 0.005 | 0.310 ± 0.001 | 0.303 ± 0.011 | 0.352 ± 0.006 |
| | pH | 6.38 ± 0.01 | 5.67 ± 0.01 | 3.42 ± 0.06 | 2.81 ± 0.30 | 2.77 ± 0.02 | 2.56 ± 0.04 | 2.420 ± 0.01 |
| ACUF731 | OD | 0.321 ± 0.005 | 0.299 ± 0.005 | 0.317 ± 0.020 | 0.364 ± 0.011 | 0.313 ± 0.005 | 0.307 ± 0.006 | 0.356 ± 0.001 |
| | pH | 6.58 ± 0.04 | 6.48 ± 0.03 | 6.14 ± 0.17 | 4.64 ± 0.00 | 3.00 ± 0.01 | 2.62 ± 0.04 | 2.22 ± 0.06 |
| ACUF551 | OD | 0.360 ± 0.006 | 0.328 ± 0.005 | 0.324 ± 0.001 | 0.327 ± 0.005 | 0.313 ± 0.021 | 0.337 ± 0.016 | 0.343 ± 0.006 |
| | pH | 6.48 ± 0.03 | 6.09 ± 0.13 | 3.54 ± 0.02 | 3.00 ± 0.00 | 2.72 ± 0.03 | 2.40 ± 0.05 | 2.16 ± 0.14 |
| **Start pH 6.5** | | **Time 0** | **Week 1** | **Week 2** | **Week 3** | **Week 4** | **Week 5** | **Week 6** |
| IPPAS P507 | OD | 0.388 ± 0.006 | 0.462 ± 0.018 | 0.564 ± 0.052 | 0.652 ± 0.068 | 0.721 ± 0.045 | 0.801 ± 0.119 | 1.402 ± 0.136 |
| | pH | 6.51 ± 0.00 | 6.40 ± 0.06 | 6.33 ± 0.04 | 6.27 ±0.17 | 6.19 ± 0.14 | 5.98 ± 0.21 | 5.68 ± 0.57 |
| ACUF769 | OD | 0.353 ± 0.001 | 0.505 ± 0.094 | 0.738 ± 0.059 | 1.282 ± 0.201 | 1.910 ± 0.144 | 3.930 ± 0.255 | 5.470 ± 0.422 |
| | pH | 6.46 ± 0.01 | 6.05 ± 0.11 | 4.76 ± 0.62 | 3.16 ± 0.11 | 2.80 ± 0.08 | 2.52 ± 0.05 | 2.33 ± 0.03 |
| ACUF722 | OD | 0.365 ± 0.007 | 0.518 ± 0.017 | 0.614 ± 0.023 | 0.827 ± 0.052 | 1.318 ± 0.003 | 1.980 ± 0.035 | 3.055 ± 0.120 |
| | pH | 6.50 ± 0.04 | 6.12 ± 0.04 | 4.26 ± 0.13 | 3.05 ± 0.04 | 2.74 ± 0.02 | 2.47 ± 0.01 | 2.25 ± 0.00 |
